## Supplemental information for "Mapping Spatial Patterns to Energetic Benefits in Groups of Flow-coupled Swimmers"

June 5, 2024

[†] Authors with equal contribution.

### Supplementary materials

#### S.1 Computational Fluid Dynamics (CFD) model

In our CFD simulations, a swimmer is modeled as a symmetric 2D Joukowsky airfoil [1]. The chord length of the airfoil is the characteristic length $L$, and the maximum thickness is $0.12L$. The airfoil undergoes pitching motion around its leading edge. Fluid-structure interactions are governed by the incompressible Navier-Stokes equations,

$$\frac{\partial \mathbf{u}}{\partial t} + \mathbf{u}\cdot\nabla\mathbf{u} = -\nabla p + \frac{1}{\mathrm{Re}}\Delta\mathbf{u}, \qquad \nabla\cdot\mathbf{u} = 0, \tag{1}$$

where $\mathbf{u}(\mathbf{x},t)$ and $p(\mathbf{x},t)$ are the velocity and pressure field, respectively. We solved for these fields numerically using immersed boundary method (IBM) that handles the two way coupled fluid structure interaction [2–7].

The immersed boundary formulation involves an Eulerian descriptions of the flow field and a Lagrangian description of the immersed swimmers, modeled as Joukowsky airfoils. The boundary condition is mapped to a body force exerted on the fluid. The Lagrangian and Eulerian variables are correlated by the Dirac delta function, which is smoothed during discretization. Here, we used the implementation developed by the group of Professor Boyce Griffith, IBAMR [8], which has long been used to solve problems such as blood flow in heart [2, 9], water entry/exit problems [10], fish's swimming [11–13], insect's flight [14, 15], flexible propulsors [16–18], self propulsion of pitching/heaving airfoil [18, 19], and fish schooling [19, 20]. This implementation is based on an adaptive mesh, which enables us to accurately simulate self propulsion and reach steady state in a large computational domain with a reasonable computational cost.

The computational domain is a rectangle of dimensions $80L \times 20L$, with periodic boundary conditions on the computational domain and no-slip boundary condition on the surface of airfoils. The initial location of the first swimmer is $12L$ away from the right boundary in streamwise direction. The initial distance between the two swimmers $d(t = 0)$ ranges from $1.5L$ to $4L$ to ensure we access different equilibria the emerge in these pairwise formations. The coarsest Eulerian mesh is a uniform $500 \times 125$ Cartesian grid. The computational domain close to the airfoils and their wake are refined. There are $3$ layers of refinement mesh, and the refinement ratio for each layer is $4$. The simulation timestep is adaptive, with the maximum timestep $\Delta t_{\max} = 2.5 \times 10^{-3}$.

The hydrodynamic forces $F_x$, $F_y$ and moment $M$ acting on each swimmer are calculated by the integrating, over the surface of that swimmer, the traction force $\boldsymbol{\sigma} \cdot \mathbf{n}$ and moment $\mathbf{x} \times (\boldsymbol{\sigma} \cdot \mathbf{n})$, where $\boldsymbol{\sigma} = -p\mathbf{I} + \mu(\nabla \mathbf{u} + \nabla \mathbf{u}^T)$ is the fluid stress tensor, $\mathbf{x}$ denotes positions on the surface of that swimmer and $\mathbf{n}$ the unit normal to that airfoil into the fluid. The hydrodynamic forces and moment are used to solve the equations of motion (1) for each swimmer and obtain the active moment $M_a$ in (6) needed to evaluate the hydrodynamic power in (7) that is expended by each swimmer.

### S.2 Vortex sheet (VS) model

In the vortex sheet model, each swimmer is modeled as a rigid plate of length $L$, and it is approximated by a bound vortex sheet, denoted by $l_b$, whose strength ensures that no fluid flows through the rigid plate, and the separated shear layer is approximated by a free regularized vortex sheet $l_w$ at the trailing edge of the swimmer. The total shed circulation $\Gamma$ in the vortex sheet is determined so as to satisfy the Kutta condition at the trailing edge, which is given in terms of the tangential velocity components above and below the bound sheet and ensures that the pressure jump across the sheet vanishes at the trailing edge. To express these concepts mathematically, it is convenient to introduce the complex notation, such that $z = x + \mathrm{i}y$, where $\mathrm{i} = \sqrt{-1}$. For more details on the vortex sheet model and our implementation of it, we refer the readers to appendix A of [21], as well as to [22–30].

The pressure difference across the infinitely thin swimmers $n[p]_\mp$ is given by integrating the balance of momentum equation for inviscid planar flow along a closed contour containing the vortex sheet and trailing edge. Here, $n = n_x + \mathrm{i}n_y$ is the unit normal, in complex notation, and $[p]_\mp$ is the jump in pressure across the swimmer. The hydrodynamic force $F = \int_{l_b} n[p]_\mp ds$ acting on the swimmer is given by,

$$\int_{l_b} n[p]_\mp ds = F_x + \mathrm{i}F_y = -F\sin\theta + \mathrm{i}F\cos\theta. \tag{2}$$

The hydrodynamic moment $M$ acting on the swimmer about its leading edge is given by

$$M = \mathrm{Re}\left[\int_{l_b} \mathrm{i}\overline{n}(z_{\mathrm{l.e.}} - z_b)[p]_\mp ds\right], \tag{3}$$

where $z_{\mathrm{l.e.}}$ is position of the leading edge $s = 0$.

We introduce a drag force $D$ that emulates the effect of skin friction due to fluid viscosity in the context of the vortex sheet model [21, 31]. Namely, following [21, 31], we write the drag force for a swimming plate

$$D = -C_d(\overline{U}_+^{3/2} + \overline{U}_-^{3/2}), \tag{4}$$

where $C_d$ is a drag coefficient and $\overline{U}_\pm$ are the spatially-averaged tangential fluid velocities on the upper and lower side of the plate, respectively, relative to the swimming velocity $U$,

$$\overline{U}_\pm(t) = \frac{1}{2l}\int_{-l}^{l} u_\pm(s,t)ds - U, \tag{5}$$

where $u_\mp(s,t)$ denotes the tangential slip velocities on both sides of the plate. We estimate $C_d$ to be approximately $0.05$ in the experiments of [32].

Additionally, following [21, 28–30], we emulate the effect of viscosity on the wake itself by allowing the shed vortex sheets to decay gradually by dissipating each incremental point vortex after a finite time $\tau_{\text{diss}}$ from the time it is shed into the fluid. Larger $\tau_{\text{diss}}$ implies that the incrementally-shed vorticity along the vortex sheet stays in the fluid for longer times, mimicking the effect of lower fluid viscosity. We studied the effect of dissipation time in Figs 1.S2 and 1.S3, and used $\tau_{\text{diss}} = 2.45T$ in the rest of this paper for the sake of computational cost.

Pitching motions are produced by an active moment $M_{\text{a}}$ imposed by the swimmer on the surrounding fluid about the leading edge. The value of $M_{\text{a}}$ is obtained from the balance of angular momentum about the swimmer's leading edge (l.e.),

$$I\ddot{\theta} - \text{Im}[m(\dot{x} + \text{i}\dot{y})w_{\text{l.e.}}] = M + M_{\text{a}}, \tag{6}$$

In the VS model, $I = mL^2/3$ is the swimmer's moment of inertia about the leading edge, $w_{\text{l.e.}}$ is the swimmer's velocity at the leading edge (in complex form), and $M$ is the hydrodynamic moment about the leading edge given in (3).

The hydrodynamic power $P$ expended by the flapping swimmer to maintain its pitching motion is given by

$$P = M_{\text{a}}\dot{\theta}. \tag{7}$$

### S.3 Time-delay particle model

We supplemented these CFD and VS models by studying pairwise interactions in the context of the minimal particle model used in [33, 34]. This particle model was designed for inline swimmers. Here, we modify it slightly to account for lateral offset $\ell$ between the swimmers as shown in Fig 5.S1. In a nutshell, the model assumes that the leader leaves behind a vertical wake speed equal to the leader's flapping speed at the tail. The speed of the 'wake' that is left behind decays exponentially in time, as an approximation for viscous dissipation. Details of the model can be found in the Supplementary Information of [34].

In this model, each swimmer is a particle of mass per unit depth $m$ undergoing vertical oscillations such that $y_1 = a\sin(2\pi ft)$ and $y = a\sin(2\pi ft - \phi)$, where $a = L\sin A$ is the tailbeat amplitude of the pitching airfoil. In this model, each particle is assumed to experience a thrust force $F_j$ that is proportional to the square of its vertical velocity relative to the surrounding fluid, and a drag force $D_j$ relative to its relative horizontal speed. Namely,

$$m\ddot{x}_j = -F_j + D_j, \qquad j = 1, 2, \tag{8}$$

where

$$F_j = \rho L C_T(\dot{y}_j - u_y(x_j, y_j))^2, \qquad D_j = \rho L C_D(\dot{x}_j - u_x(x_j, y_j))^2. \tag{9}$$

Here, $u_x$ and $u_y$ are the $x$- and $y$-components of the fluid velocity, $C_T, C_D$ are constant thrust and drag coefficients. Since the leader swims into quiescent fluid, we assume that $u_y(x_1, y_1) = 0$. The follower swims into the wake of the leader, which we assume to have zero horizontal velocity ($u_x = 0$) and vertical velocity that decays exponentially both in time and in the lateral direction,

$$u_y(x_2, y_2) = \dot{y}_1(t - \Delta t)e^{-\Delta t/\tau}e^{-|\ell/h|^p}, \tag{10}$$

where $\Delta t$ is the delay time between the leader and follower, that is, the time past since the leader passed by the follower's current location: $x_1(t - \Delta t) = x_2(t)$ We added the last term in (10) to consider decay in the lateral direction $\ell$; to estimate the parameters $p$ and $h$, we used a best curve fit to the data of period-average velocity magnitude versus lateral distance in the wake of a single swimmer in the vortex sheet model (Fig. 5.S1). We numerically integrated (9) and solved for the motion of the follower. At steady state, we computed the separation distance $d = x_2 - x_1$ between the pair acting on the follower for a range of $\phi \in [0, 2\pi]$ and $\ell \in [-L, L]$ (Fig. 5.S1). The parameter values are chosen to be consistent with the experiments of Newbolt *et al.* (provided in Table S1 of SI in [34]). Specifically, $\rho = 1\,$g/cm$^3$, $L = 4\,$cm, $m = 5.3\,$g/cm, $C_D = 0.25$, $C_T = 0.96$, $\tau = 0.5\,$sec.

#### S.4 Flow agreement and thrust parameters

The flow agreement parameter field is defined

$$\mathbb{V}(x, y; \phi) = \frac{1}{\int_t^{t+T} \mathbf{v} \cdot \mathbf{v}\, \mathrm{d}t'} \left[ \int_t^{t+T} \mathbf{v} \cdot \mathbf{u}\, \mathrm{d}t' \right], \tag{11}$$

where $\mathbf{u}(x, y, t)$ is the flow field of an oncoming wake and $\mathbf{v}(t; \phi)$ is flapping velocity of a virtual particle located at a location $(x, y)$ in the oncoming wake. The thrust parameter field is defined as

$$\mathbb{T}(x, y, \phi) = -\frac{1}{\int_t^{t+T} |\mathbf{v} \cdot \mathbf{e}_y|^2\, \mathrm{d}t'} \left[ \int_t^{t+T} |(\mathbf{v} - \mathbf{u}) \cdot \mathbf{e}_y|^2\, \mathrm{d}t' \right]. \tag{12}$$

To further illustrate the meaning of these parameters, we considered an ideal scenario, where the flow velocity is simply of the form of

$$\mathbf{u} = u_x \mathbf{e}_x + u_y \mathbf{e}_y = -U \mathbf{e}_x + 2\pi f A_u \cos(2\pi f t) \mathbf{e}_y, \tag{13}$$

where $u_y(t) = 2\pi f A_u \cos(2\pi f t)$ is the transverse fluid velocity. We placed a virtual or "ghost" swimmer heaving at a velocity $v(t; \phi) = 2\pi f A_v \cos(2\pi f t - \phi)$. Under this scenario, flow agreement parameter is

$$\mathbb{V} = \frac{A_u}{A_v} \cos\phi \tag{14}$$

and thrust parameter is

$$\mathbb{T} = A_u^2 + A_v^2 - 2A_v A_u \cos\phi \tag{15}$$

Let's now consider a balance of forces on the "ghost swimmer." The ghost swimmer is in relative equilibrium if the sum of drag forces $D$ and thrust forces $\mathbb{T}$ are zero,

$$D + \mathbb{T} = 0 \quad \Rightarrow \quad \cos\phi = \frac{A_u^2 + A_v^2 - D}{2A_v A_u} \quad \Rightarrow \quad \phi = \pm \cos^{-1} \frac{A_u^2 + A_v^2 - D}{2A_v A_u}. \tag{16}$$

Substituting back into (14), we get

$$\mathbb{V} = \frac{A_u^2 + A_v^2 - D}{2A_v^2} \tag{17}$$

According to our prediction based on the flow agreement parameter, equilibria are located at maximal flow agreement parameter, for which $\phi = 0$ and $\mathbb{V} = A_u/A_v$. This is close to the calculated values of $\phi$ and $\mathbb{V}$ when $A_u \approx A_v$ and $D \approx 0$, that is, assuming the ghost swimmer is getting a free ride with the local flow at zero thrust and drag.

To consider stability at the calculated equilibria in (16), we calculate the derivative of thrust parameter in (15) with respect to phase $\phi$ and evaluate it at the equilibrium points

$$\frac{\partial \mathbb{T}}{\partial \phi} = -2A_v A_u \sin\phi = \mp 2A_v A_u \sqrt{1 - \frac{A_u^2 + A_v^2 - D^2}{2A_v A_u}} \tag{18}$$

Thus, there is always a stable equilibrium and an unstable equilibrium, both of them are close to the local maximum of flow agreement parameter $\mathbb{V} = A_u/A_v$, which is consistent with Fig. 4 of the main text.

### S.5 Phase control

Consider a swimmer flapping at a phase $\phi$ can 'sense' or measure the agreement of its flapping motion with the local fluid velocity $\mathbf{u}$ (generated by sources other than itself) at its location (say at its midpoint), over a time span of $m$ flapping periods. The goal of the swimmer would be to adjust its current flapping phase $\phi$ to a desired phase $\Phi$ that maximizes its agreement with the local velocity

$$\Phi(t) = \underset{\phi}{\text{argmax}} \frac{\frac{1}{mT}\int_{\min(t-mT,0)}^{t} \mathbf{v}(t',\phi)\cdot\mathbf{u}(t')\,\mathrm{d}t'}{\frac{1}{mT}\int_{\min(t-mT,0)}^{t} \mathbf{v}(t',\phi)\cdot\mathbf{v}(t',\phi)\,\mathrm{d}t'}, \tag{19}$$

using a proportional phase controller inspired from [35, 36],

$$\ddot{\phi}(t) = -\gamma^2\left[\phi(t) - \Phi(t)\right] - 2\gamma\dot{\phi}(t); \tag{20}$$

Here, $\Phi(t)$ is the desired phase and $\gamma$ is a constant that determines the speed of convergence. We chose the parameters as follows: we set the number of periods $m = 2$ that describes the memory of the swimmer of the ambient fluid $\mathbf{u}$, such that the time history is two times the pitching period $2T$. We set the control gain $\gamma = 3$ to ensure that the actual phase $\phi(t)$ can reach the desired phase $\Phi(t)$ at $1\%$ of relative error within $1.5T$.

By implementing this phase controller in swimmer 4 in a group of four inline swimmers, the swimmer is able to stabilize itself in the formation as shown in Figure 10. However, this stabilization is expensive: by actively controlling its phase to stay in formation, swimmer 4 spends 100% more hydrodynamic power than swimming alone.

#### S.6 Data collection from published literature

To probe the relationship between the scaled separation distance $d/UT$ in pairs of swimmers and their flapping phase lag $\phi$, we collected data from published literature on pairs of interacting swimmers [21, 34, 36–38]; see Table S.1 and the Supplemental Excel Sheet. We excluded a few studies from this literature survey [39, 40], because we were unable to extract $d/UT$ from the data they provided. We also excluded [33] because their study involved fixed inter-swimmer distances.

We found that the relationship between phase lag $\phi$ and separation distance $d/UT$ is approximately linear $\phi/2\pi = d/UT + \beta$. The value of $\beta$ depends on how one defines the inter-swimmer separation distance. The two definitions that are commonly used in the literature are tail-to-tail or tail-to-head (gap) distance. These definitions do not change the linear scaling between $d/UT$ and $\phi/2\pi$; they only change the value of $\beta$. For the data provided in Fig. 3 of the main text, we used the definition of separation distance $d$ that results in $\beta = 0$. Not all literature provided values for $U$ or $T$. In the very few cases when this information were missing, we estimated $UT$ from the linear relationship $\phi/2\pi = d/UT$ [37].

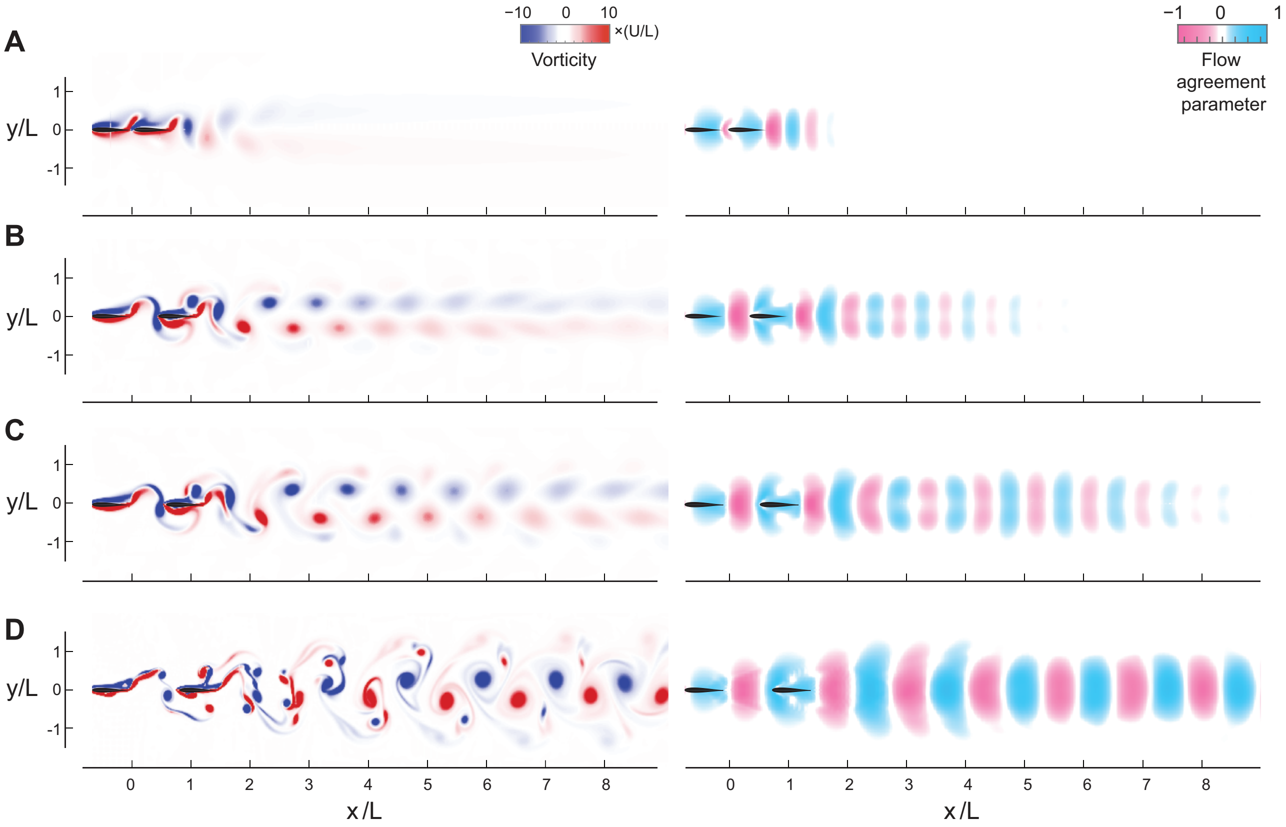


Figure 1.S1: **Pairs of swimmers in CFD simulations** Vorticity field (left column) and flow agreement parameter $\mathbb{V}$ (right column) in the wake of a pair of inline and inphase swimmers Reynolds number $\mathrm{Re}_A = 206, 308, 411, 1645$, respectively. The pitching amplitude of leader and follower is set to $A = 15°$, except in **A**, where the follower is pitching at $A = 13.5°$.

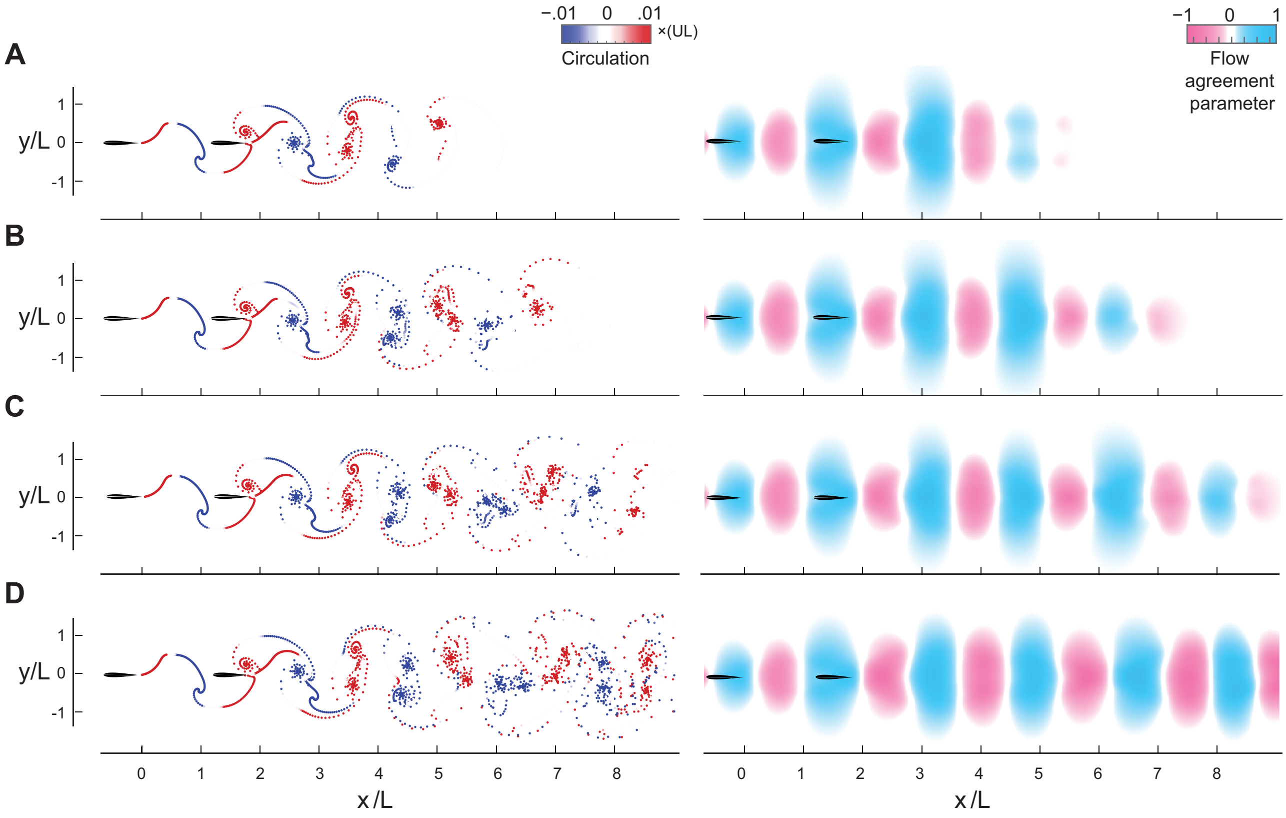


Figure 1.S2: **Pairs of swimmers in VS simulations** Snapshots of the swimmers (left column) and flow agreement parameter $\mathbb{V}$ (right column) in the wake of a pair of swimmers in the VS model for dissipation time $\tau_{\text{diss}} = 2.45T$, $3.45T$, $4.45T$, and $9.45T$, respectively. The pitching amplitude is set to $A = 15°$, and the pitching frequency to $f = 1$.

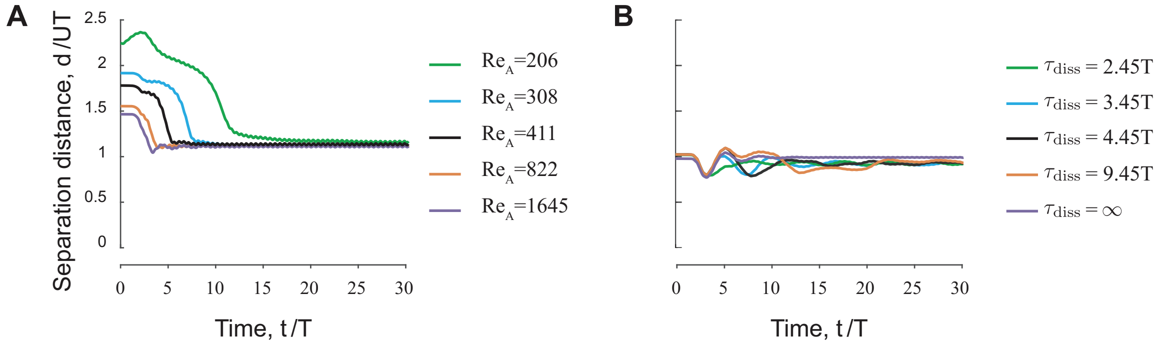


Figure 1.S3: **Influence of fluid property. A.** Separation distance versus time in a pair of swimmers in the CFD model for five values of Reynolds numbers Re$_A$ $= 206, 308, 411, 822, 1645$ (Fig. 1.S1). **B.** Separation distance versus time in a pair of swimmers in the VS model for five values of dissipation time $\tau_{\text{diss}} = 2.45T, 3.45T, 4.45T, 9.45T, \infty$ (Fig. 1.S2). Separation distance $d$ is normalized by swimming speed $U$ for each cases, respectively. In **A.**, **B.**, the swimmers stabilize near $d/UT = 1$. The pitching amplitude of leader and follower is set to $A = 15°$, except in Re$_A$ $= 206$ , where the follower is pitching at $A = 13.5°$ to avoid collision.

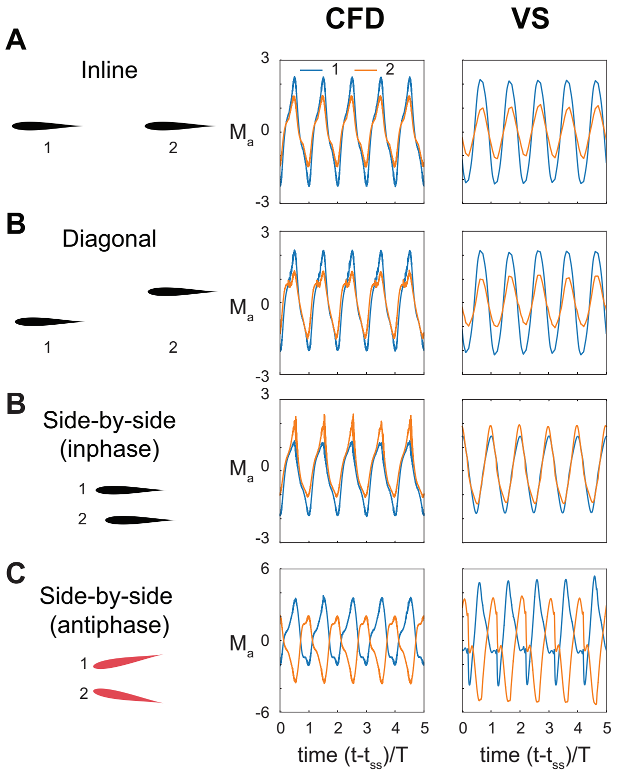


Figure 1.S4: **Hydrodynamic torque for pair of swimmers at different spatial configurations. A.** inline ($\ell = 0, \phi = 0$), **B.** diagonal ($\ell = L/4, \phi = 0$), **C.** inphase side-by-side ($\ell = L/2, \phi = 0$) and **D.** antiphase side-by-side ($\ell = L/2, \phi = \pi$) in CFD (left) and VS (right) simulations. For each simulation, we show the active torque $M_a$ exerted by the swimmers after a time $t_{ss}$, ensuring that steady state has been reached. The non-reciprocity in the effects of leader on follower in inline and diagonal configuration is apparent. In the side-by-side confirgurations, a simple shift of the data in the inphase flapping case and a simple mirror symmetry in the antiphase flapping case would show that flow coupling is reciprocal.

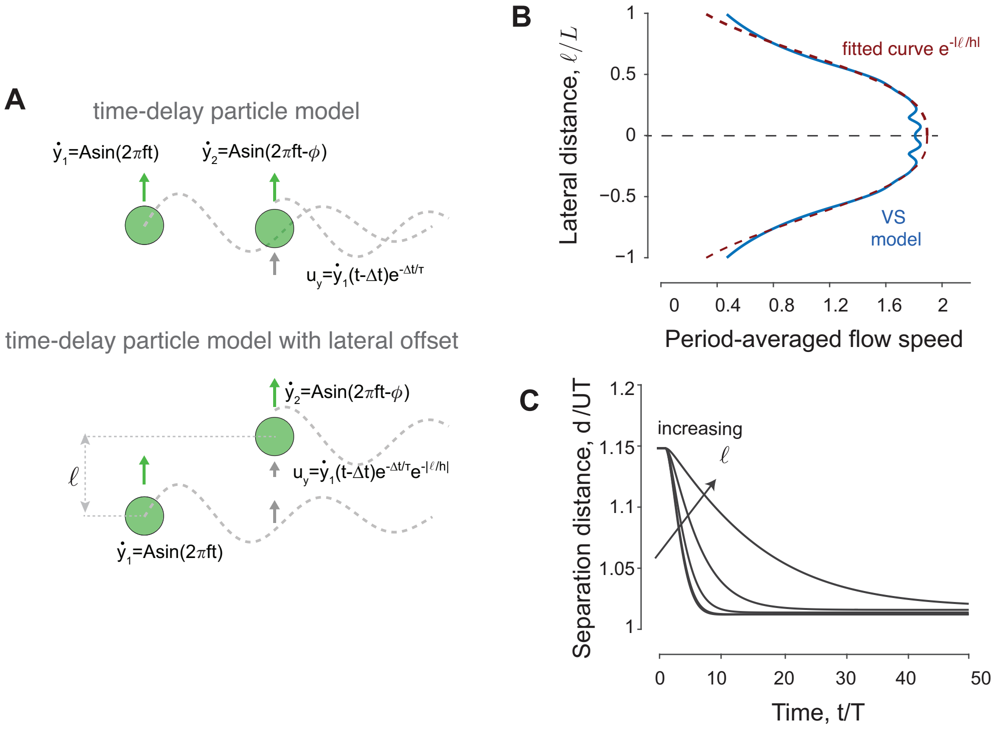


Figure 5.S1: **Time-delay particle model and stability of swimmer. A.** Schematics of time-delayed particle model [32, 34] and its extension to laterally-offset swimmers. Each swimmer generates hydrodynamic thrust via oscillating vertically, which also leaves a wake behind it. The follower swimmer interacts with the wake of the leader. **B.** For an inphase pair, starting at initial distance $d/UT = 1.15$, we incrementally increase the lateral offset from $\ell = 0$ to $\ell = L$. (inset) Lateral decay of flow speed in the wake of a solitary swimmer in the vortex sheet model (blue line) and the fitted exponential curve (red line). The lateral exponential decay in the time-delay particle model takes the form $\exp\left(-|\ell/1.6|^{2.73}\right)$. **C.** The hydrodynamic force $\langle F_2 \rangle$ acting on the follower as a function of the corresponding distance from the equilibrium. Due to the decay of the leader's wake in the lateral direction (see inset in A), the hydrodynamic force $\langle F_2 \rangle$ experienced by the follower decreases in magnitude at increasing lateral offset $\ell$. Orange lines show the linear change in force at the corresponding equilibria. The slope $\delta F/\delta d$ is a measure of linear stability. Negative slopes imply stable formations for all $\ell \leq L$, but these slopes become more shallow as we increase $\ell$.

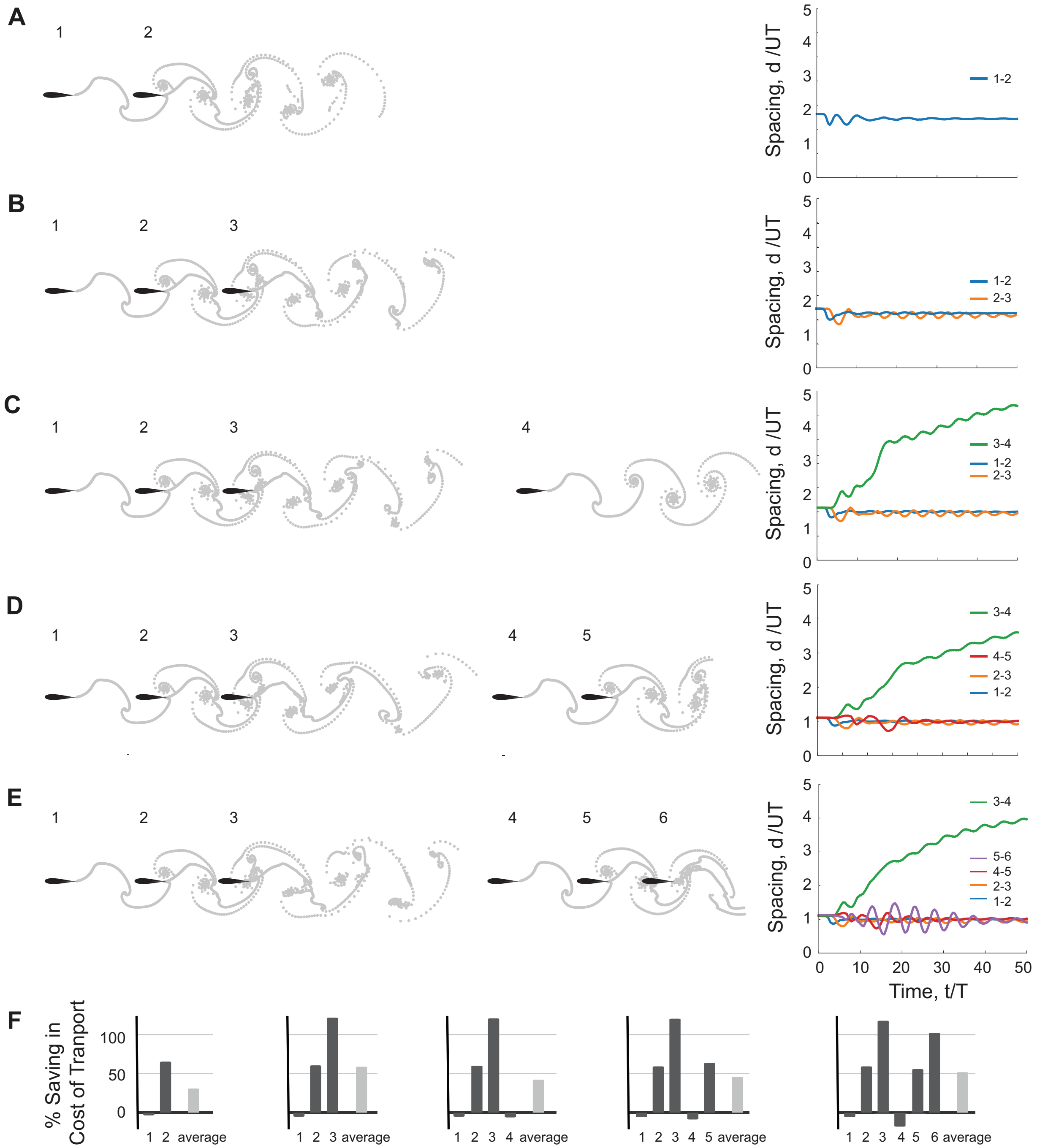


Figure 7.S1: **Inline formations.** Snapshots of inline formations composed of 2, 3, 4, 5 and 6 swimmers in VS simulations at steady state; time-evolution of pairwise distances is shown on the right. **A** and **B.** Formations composed of 2 or three swimmers are stable with at consecutive spacing $d/UT = 1$. **C.** For a trail of 4 swimmers, the group splits into a leading subgroup of 3 swimmers while the fourth swimmer separates from the rest. **D** and **E.** For formation of 5 or 6 swimmers, the group splits into a leading subgroup of 3 swimmers and another subgroup containing the remaining 2 or 3 swimmers. **F.** reports recent savings in COT for each swimmer and the average of the whole group.

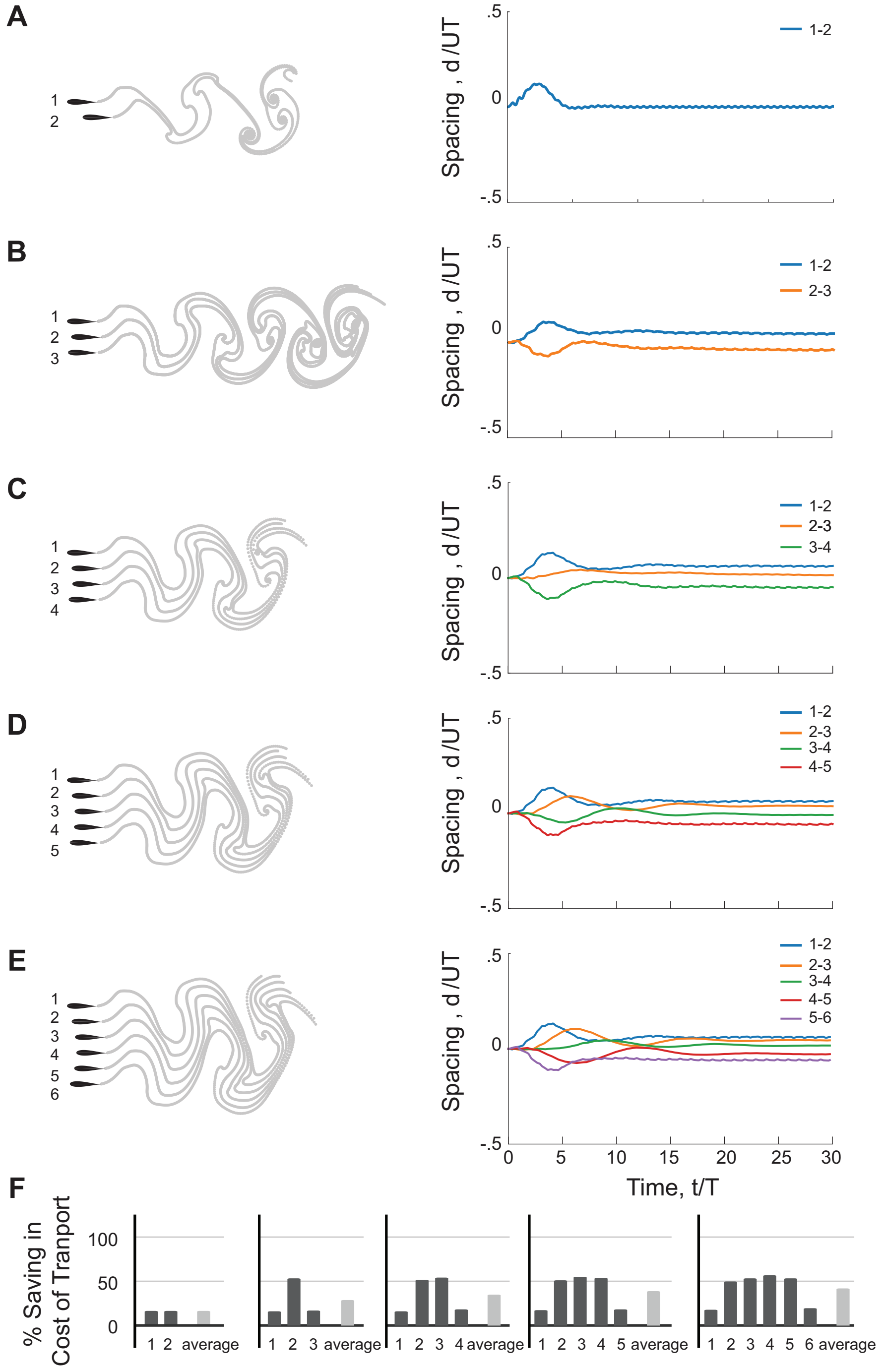


Figure 7.S2: **Side-by-side formations.** Vortex sheet simulation of side-by-side formations with 2, 3, 4, 5 and 6 swimmers, from top to bottom, respectively. On right hand side, we report pairwise spacing between them. In all of the groups the formations are stable and the distances between every pair are close to zero. **F.** reports recent savings in COT for each swimmer and the average of the whole group.

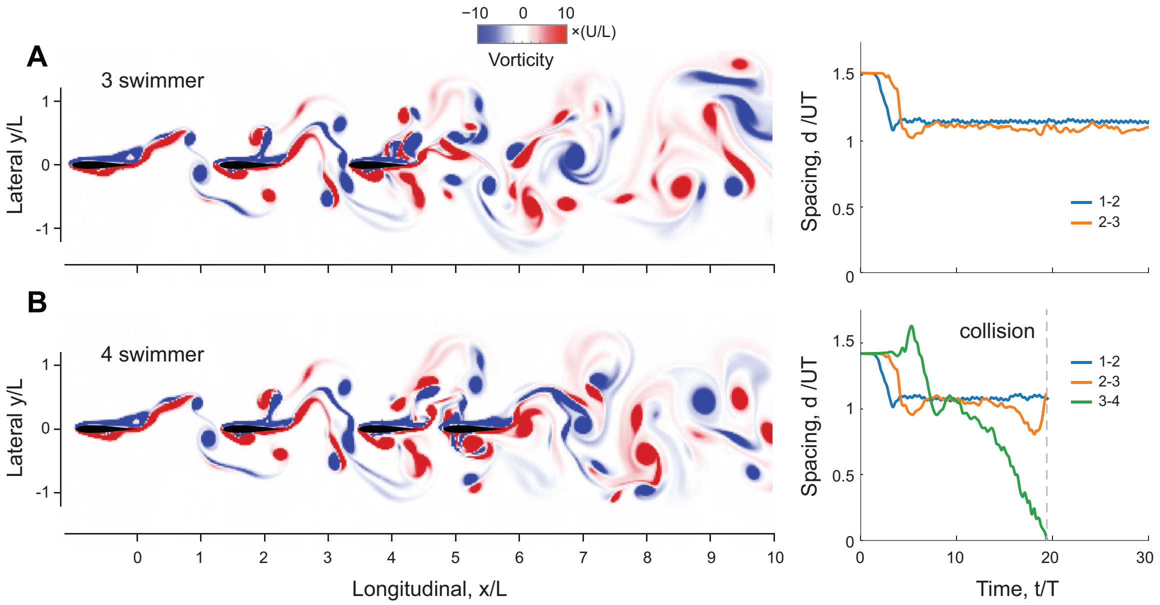


Figure 9.S1: **CFD simulation of larger inline schools.** CFD simulation of inline formations with 3 and 4swimmers at Re $= 1645$. Vorticity field is shown on the left hind side. On right hand side, we report pairwise spacing between them. The pitching amplitude of leader and follower is set to $A = 15°$.

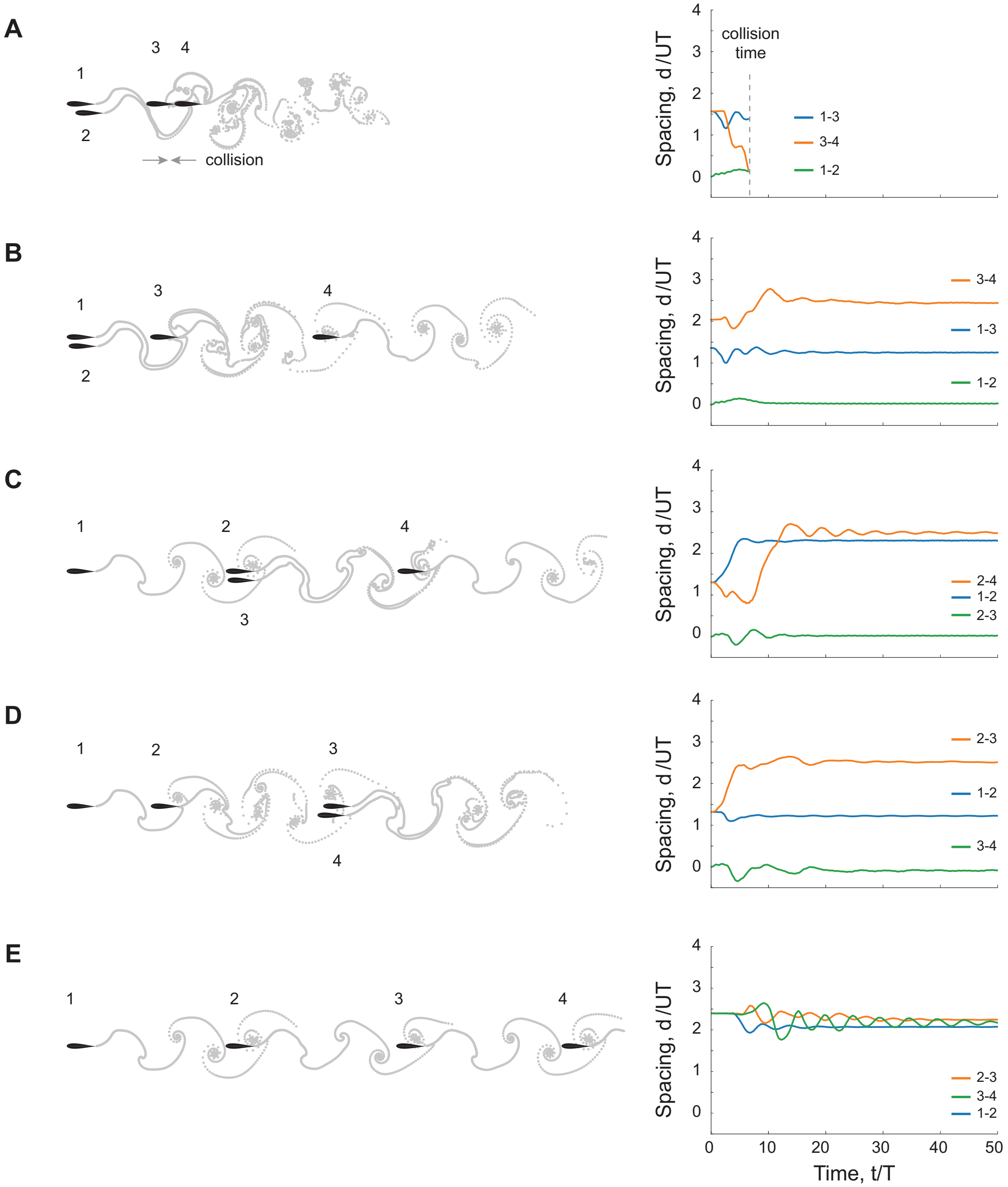


Figure 10.S1: **Alternative formations of four swimmers** Vortex sheet simulation of four swimmers with alternative formations. On right hand side, we report pairwise spacing between them. The lateral distance is $\ell = 0.25L$. **A.** Two leading swimmers swim side by side. The third and forth swimmer collide to each other. **B.** The same configuration as in **A.** with larger initial distance between the third and forth swimmer. They form a stable school. The forth swimmer stays at the second equilibrium behind the third swimmer. **C.** An additional swimmer is placed side by side to the second swimmer in a school of three inline swimmers. The second and third swimmer stays at the second equilibrium from the first swimmer, and the forth swimmer stays at the second equilibrium from the second and third swimmer. **D.** An additional swimmer is placed side by side to the third swimmer in a school of three inline swimmers. The last two swimmers stay at the second equilibrium from the second swimmer. **E.** Four inline inphase swimmers when initially placed close to the second equilibrium. Power saving per swimmer is reported in Fig. 10.

[1]

Table S.1: Data collection from published literature. We calculated the Reynolds numbers based on swimming speed ($\mathrm{Re}_U = \rho U L/\mu$) and flapping velocity ($\mathrm{Re}_A = 2\pi\rho A f L/\mu$), where $f = 1/T$ is the flapping frequency. Missing values are either not applicable or not available.

| **reference** | **study type** | **system** | **distance** | **phase** | $\mathrm{Re}_U$ | $\mathrm{Re}_A$ |
|---|---|---|---|---|---|---|
| Kim *et al.* 2010 | Numerical | Flapping flags | head-to-head | $\phi \in [0, 2\pi]$ | $200$ - $400$ | - |
| Zhu *et al.* 2014 | Numerical | Heaving flexible foils | - | $\phi \in [0, \pi]$ | $500$ | $200$ |
| Ramananarivo *et al.* 2016 | Experimental | Heaving foils | tail-to-head | $\phi = 0$ | $10^3$- $10^4$ | $10^2$- $10^3$ |
| Peng *et al.* 2018 | Numercial | Heaving flexible foils | tail-to-head | $\phi = 0$ | $509$ | $200$ |
| Newbolt *et al.* 2019 | Experimental | Heaving foils | tail-to-head | $\phi \in [0, 2\pi]$ | $10^3$ - $10^4$ | $10^2$ - $10^3$ |
| Li *et al.* 2020 | Experimental | Goldfish | head-to-head | $\phi \in [0, 2\pi]$ | $10^5$ | $10^4$ |
| Kurt *et al.* 2021 | Experimental | Pitching foils | tail-to-head | $\phi \in [0, 2\pi]$ | $9950$ | $18850$ |
| Heydari & Kanso 2021 | Numerical | Heaving plates | tail-to-head | $\phi = 0$ | - | - |
| | | Pitching plates | head-to-head | $\phi = 0$ | - | - |
| Arranz *et al.* 2022 | Numerical | Heaving flexible foils (3D) | tail-to-head | $\phi \in [0, 2\pi]$ | $176$ | $200$ |
| Thandiackal & Lauder 2023 | Experimental | Fish-foil interactions | tail-to-head | $\phi \in [0, 2\pi]$ | $20 \cdot 10^3$ (foil)<br>$40 \cdot 10^3$ (fish) | $5 \cdot 10^3$ (foil) |
| * Becker *et al.* 2015 | Experimental | Heaving foils | tail-to-head | $\phi = 0, \pi$ | - | $10^2$ - $10^4$ |
| * Park *et al.* 2018 | Numerical | Heaving flexible foils | - | $\phi \in [0, 2\pi]$ | $60$ - $1100$ | $100$ - $1200$ |
| * Dai *et al.* 2018 | Numerical | Pitching + heaving foils | - | $\phi = 0, \pi$ | $440$ | $600$ |

[1] * data not included in Fig. 3.
